# Highly-sensitive reporter cell line for detection of interferon types I-III and their neutralization by antibodies

**DOI:** 10.1101/2024.06.18.599537

**Authors:** Kevin Groen, Roger Kuratli, Andri Vasou, Michael Huber, David J. Hughes, Benjamin G. Hale

## Abstract

Interferons (IFNs) are a critical component of innate immune defenses and limit viral disease severity. To advance studies on IFNs and their neutralization by pathogenic autoantibodies, we generated a Renilla luciferase-based reporter cell line capable of detecting the activities of IFN-Is, IFN-II, and IFN-IIIs. The reporter cell line exhibits a 125-to 2000-fold higher sensitivity to IFNs than a commonly-used alternative biological reporter system, and allows for a rapid and simple live-cell workflow for detecting low titer amounts of neutralizing anti-IFN antibodies.

## Main text

The human interferon (IFN) system constitutes a critical component of innate immunity. Upon detection of infection, cells can initiate the production and secretion of several IFN cytokine types (IFN-I, IFN-II, and IFN-III). These IFNs act to trigger expression of a myriad of IFN-stimulated genes (ISGs), causing cells to enter an antiviral state. IFN-Is (e.g. IFNα, IFNβ, and IFNω) and IFN-IIIs (e.g. IFNλ1-4) are most critical to this defense, and signal via type-specific receptors expressed on the surface of either a broad range of cells (IFN-I receptor) or a tissue-restricted set of cells (IFN-III receptor)^1^. Deficiencies in the human IFN system can seriously exacerbate the severity of infections^2^, and a functional IFN deficiency caused by autoantibodies (autoAbs) neutralizing IFN-Is (mostly IFNα and IFNω) promotes the replication of several viral pathogens and thereby exacerbates disease progression^3–7^. AutoAbs against IFN-II (IFN*γ*) and IFN-IIIs have also been described, but their precise contributions to viral disease are currently unclear^8,9^. Identification of individuals harboring neutralizing anti-IFN autoAbs is therefore critical to understand susceptibility to severe infections, and requires highly-sensitive and robust assays.

Previously, neutralizing anti-IFN-I autoAbs have been detected by transfecting 293T cells with plasmids expressing Firefly luciferase under control of an ISG (*MX1*) promoter, followed by stimulation with recombinant IFNs pre-incubated with patient plasmas. The ability of patient autoAbs to neutralize IFNs is then inferred from reduced levels of IFN-induced Firefly luciferase activity 16-24h later^6,10^. However, this system has several caveats: (i) transfection of cells adds 24-48h to assay duration and increases inter-batch variability (controlled by co-transfection of a constitutively-expressed reporter); (ii) relatively high IFN amounts are required to induce signaling, limiting detection of low titers of neutralizing autoAbs; (iii) peak IFN-induced Firefly luciferase requires 24h, extending assay duration; and (iv) 293T cells do not respond to IFN-IIIs^11^, prohibiting their use to study anti-IFN-III autoAbs. To overcome these caveats, we developed a highly-sensitive reporter cell line for detection of IFN-Is, IFN-II, and IFN-IIIs. The cell line is based on A549s, which respond efficiently to all IFN types^12^. A549 cells were generated with an integrated gene cassette for monocistronic expression of both Renilla luciferase and puromycin resistance downstream of the human *ISG15* promoter (prISG15) (**Fig. 1A**)^13^. The *ISG15* promoter was chosen for its inducibility by IFN-Is, IFN-II, and IFN-IIIs, as well as the observation that IFN-stimulated *ISG15* levels are maintained for periods of up to 48h^14^. Renilla luciferase was chosen as the reporter enzyme due to its inherently high signal-to-noise ratio and the availability of a live-cell substrate, which allows for uncomplicated temporal monitoring of luciferase activity. To enhance sensitivity, Renilla luciferase was expressed with a C-terminal PEST (peptide rich in proline, glutamate, serine, and threonine) sequence that promotes protein degradation, thereby reducing any background luciferase from leaky promoter activity^15^. Following generation of an initial polyclonal set of cells, single-cell clones were stimulated with 1,000 international units (IU)/mL of IFNα2 or IFNλ1 to induce prISG15-mediated Renilla luciferase activity. Among 66 tested clones, one (which had particularly high Renilla induction levels following stimulation) was selected for use (**Fig. 1B**). This clone was named the A549 IFN-reporter (AIR) cell line.

**Figure 1.**
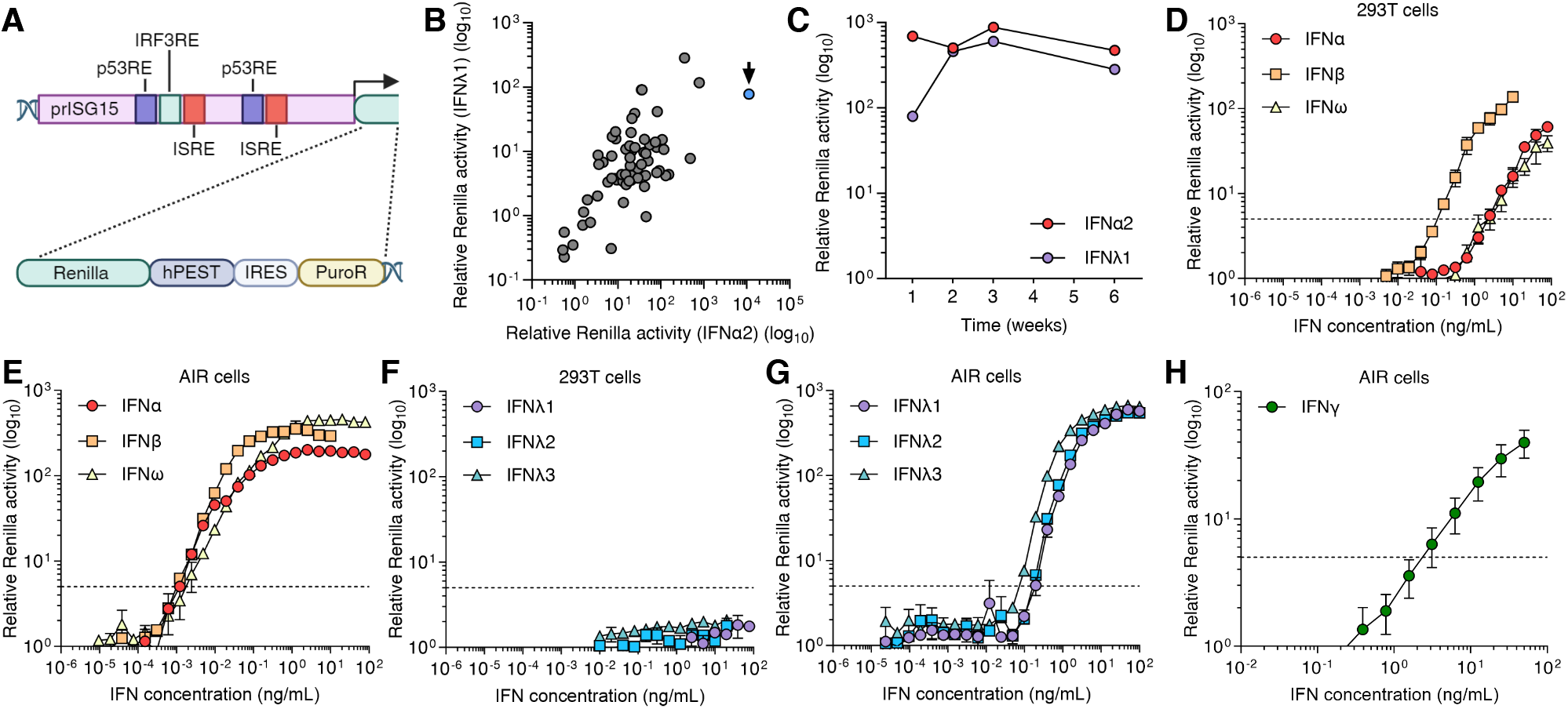
Generation of a highly-sensitive A549-IFN-reporter (AIR) cell line. (**A**) Schematic representation of the gene cassette used to transduce A549 cells, adapted from ^13^ and created with BioRender.com. Abbreviations: p53RE, p53-responsive element; IRF3RE, IRF3-responsive element; ISRE, IFN-stimulated response element; PuroR, puromycin resistance gene; IRES, internal ribosome entry site. (**B**) Renilla luciferase activity following stimulation of 66 transduced cell clones with IFNα2 or IFNλ1 (1000 IU/mL) for 24h. Arrow indicates the clone (AIR) selected for further characterization. (**C**) Stability of IFNα2 and IFNλ1 responsiveness (1000 IU/mL, 24h) in AIR cells after passaging over 6 weeks. (**D-G**) Sensitivity of a standard 293T-based assay (D, F) and AIR cell assay (E, G) to stimulation with the indicated concentrations of IFN-Is (α2, β, ω; 24h; D-E) or IFN-IIIs (λ1-3; 24 h; F-G). (**H**) Sensitivity of AIR cells to stimulation with the indicated concentrations of IFN-II (γ, 24h). Dashed lines in D-H indicate 5-fold luciferase induction compared to non-stimulated cells. Mean values from n=3 replicates are shown. Error bars represent standard deviations.

First, we confirmed stable responsiveness of AIR cells to IFNα2 or IFNλ1 following passaging over six weeks (**Fig. 1C**). We then compared the IFN-sensitivities of AIR cell assays with those of the standard 293T transfection-based dual luciferase system. Both reporter systems were stimulated with a concentration range of IFN-Is (IFNα2, IFNβ, and IFNω) or IFN-IIIs (IFNλ1-3), and the respective IFN-induced luciferase activities were determined 24h later. The sensitivity to each IFN was defined as the lowest IFN dose that induced activity >5-fold above the non-stimulated condition. With this criterion, the 293T system was able to detect 2.5 ng/mL IFNα2, 150 pg/mL IFNβ, and 2.5 ng/mL IFNω (**Fig. 1D**). By comparison, the AIR system was able to detect levels of IFN as low as 1.2 pg/mL IFNα2 or IFNβ, and 2.4 pg/mL for IFNω (**Fig. 1E**). In addition, while 293T cells did not respond to stimulation with IFN-IIIs (**Fig. 1F**), AIR cells responded to as little as 195 pg/mL IFNλ1 or IFNλ2, and 97.5 pg/mL IFNλ3 (**Fig. 1G**). Since *ISG15* is upregulated by IFNγ stimulation of A549 cells^14^, we also assessed AIR cell sensitivity to IFNγ. Indeed, AIR cells responded to as low as 3.1 ng/mL IFNγ (**Fig. 1H**). Thus, AIR cells display a 125-to 2000-fold higher sensitivity to IFNs than 293Ts, and universally react to all three IFN types.

Next, we compared IFN-induced luciferase kinetics and stability between the AIR cell system and the 293T-based system. Using a single dose of each IFN (10 ng/mL IFNα or IFNω, 1.21 ng/mL IFNβ, and 50 ng/mL IFNλ1-3), the earliest robust induction of luciferase (>5-fold over non-stimulated) in 293Ts occurred at 6-8h post-stimulation, and only peaked at 24-30h (**Fig. 2A**). In contrast, the earliest luciferase signal in AIR cells was already detectable after 1h of stimulation, and peaked after 8h (**Fig. 2B**). Importantly, this signal remained stable in AIR cells for 30h, thereby providing a rapidly-induced and long-lasting experimental window. Compared to the lengthy transfection steps, slow reactivity, and cell lysis requirements in transfection-based 293T assays, the speed and sensitivity of AIR cell assays allows a faster and simpler workflow for determining IFN activity.

**Figure 2.**
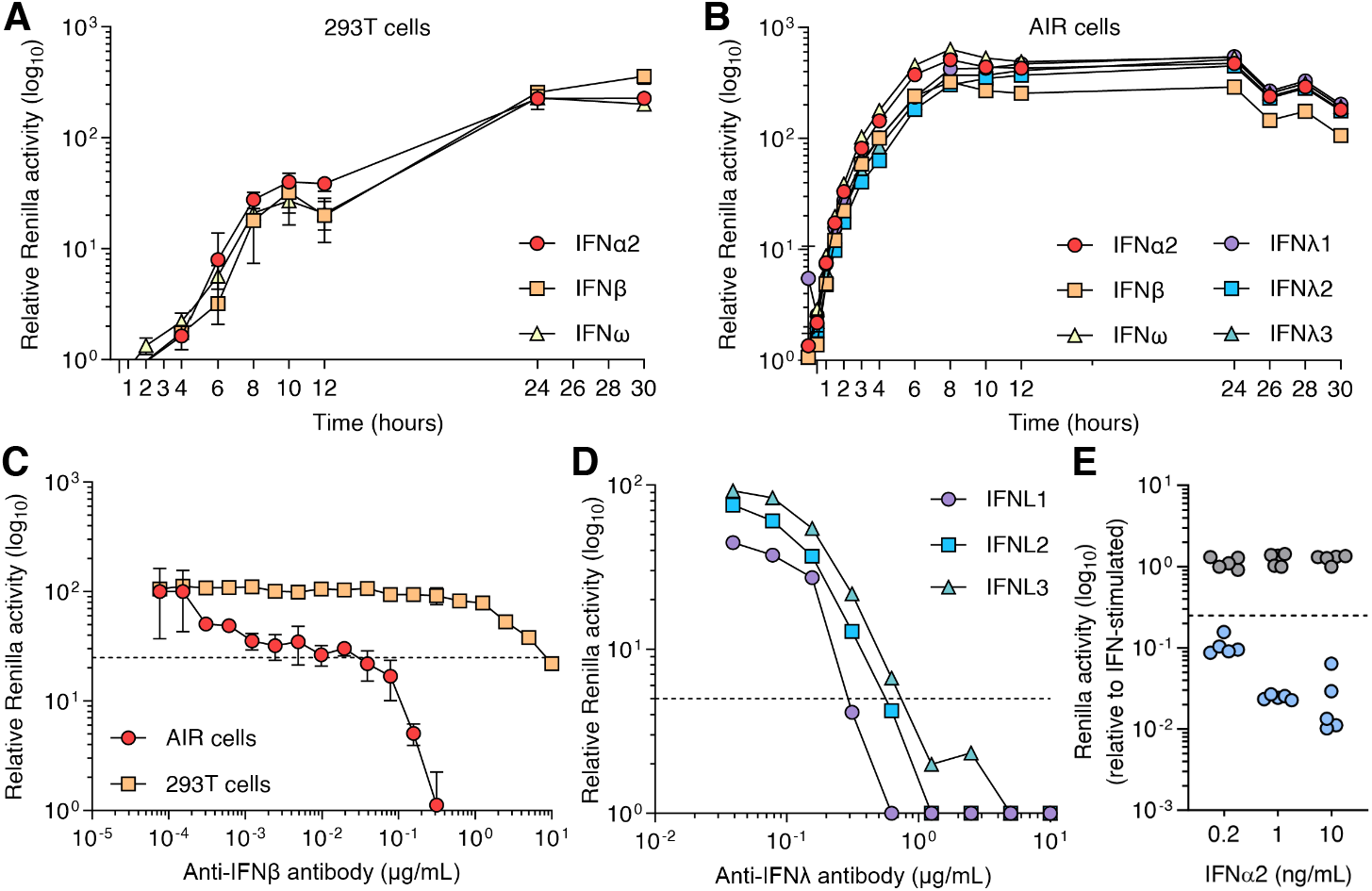
Characterization of AIR cells and their application to detect neutralizing anti-IFN antibodies. (**A-B**) Renilla luciferase induction kinetics over 30h following stimulation of 293T cells (A) or AIR cells (B) with IFN-Is (IFNα2 and IFNω: 10 ng/mL, IFNβ: 1.21 ng/mL) or IFN-IIIs (λ1-3; 50 ng/mL). (**C**) Sensitivity analysis of 293T cells and AIR cells to detect neutralization of IFNβ by an anti-IFNβ antibody. The indicated concentration of antibody was incubated with 200 pg/mL IFNβ or 3.6 pg/mL IFNβ prior to stimulation of 293T or AIR cells, respectively. (**D**) Sensitivity analysis of AIR cells to detect neutralization of IFN-IIIs by their respective antibodies. The indicated concentration of antibody was incubated with 10 ng/mL IFNλ1, IFNλ2, or IFNλ3 prior to stimulation of AIR cells. (**E**) Use of AIR cells to determine the neutralization of IFNα2 at the indicated concentration by human plasmas. Blue: plasmas with neutralizing IFNα activity. Gray: negative control plasmas. The dashed lines in C-E indicate the 75% neutralization threshold. For A-C, mean values from n=3 replicates are shown and error bars represent standard deviations. For D-E, data are representative of at least n=2 similar experiments.

Finally, we assessed the sensitivity of AIR cell assays to detect neutralizing anti-IFN antibodies. To test this, we initially used the lowest amount of IFN-I that resulted in a clear 10-fold induction of luciferase signal over non-stimulated cells (200 pg/mL IFNβ for 293T cells and 3.6 pg/mL IFNβ for AIR cells). These IFNβ amounts were incubated with a dilution series of a commercially-available neutralizing anti-IFNβ antibody for 1h prior to stimulation of each cell system. With a neutralization threshold of 75% reduction, the 293T assay was able to detect 10 µg/mL of neutralizing antibody, while AIR cell assays had a 250-fold improved sensitivity, detecting neutralization with only 39 ng/mL antibody (**Fig. 2C**). Similar assays demonstrated that AIR cells can be used to detect neutralizing anti-IFN-III antibodies in a dose-dependent manner (**Fig. 2D**). Finally, we selected five previously characterized human plasmas that neutralize IFNα activity (and five negative plasmas)^10^, and demonstrated the ability of AIR cells to detect neutralizing activity in such specimens (**Fig. 2E**).

In summary, these data demonstrate that the high sensitivity of AIR cells to respond rapidly to very low amounts of all IFN types permits their application to detect rare and low titer neutralizing anti-IFN autoAbs.

## Material and methods

### Cells and media

293T and A549 cells were originally from ATCC, and were cultured in Dulbecco’s Modified Eagle’s Medium (DMEM; Thermo Fisher Scientific) supplemented with 10% fetal bovine serum (FBS) and 100 U/mL penicillin and streptomycin (Gibco Life Technologies; 15140-122). Cells were grown at 37°C with 5% CO_2_.

### Human plasma samples and ethics

Anonymized human plasma samples previously categorized as positive (n=5) or negative (n=5) for anti-IFNα autoAbs^10^ were used to validate assays. The samples were originally derived from specimens stored in the Zurich center biobank of the Swiss HIV Cohort study (SHCS,^16^) which covers their use in the current work. Detailed information on the study is available at http://www.shcs.ch. The SHCS has been approved by the ethics committees of all participating institutions (Kantonale Ethikkommission Bern, Ethikkommission des Kantons St. Gallen, Comite Departemental d’Ethique des Specialites Medicales et de Medicine Communataire et de Premier Recours, Kantonale Ethikkommission Zürich, Repubblica et Cantone Ticino–Comitato Ethico Cantonale, Commission Cantonale d’Étique de la Recherche sur l’Être Humain, Ethikkommission beider Basel), and written informed consent has been obtained from all participants.

### Cloning of plasmids

Plasmid prISG15-eGFP-IRES-Puro has been described previously^17^. For cloning of plasmid prISG15-Renilla-hPEST-IRES-Puro, the Renilla luciferase sequence was PCR amplified from plasmid pRL-TK-Renilla (Promega, E2241) and the hPEST sequence was PCR amplified from plasmid pcDNA3.1(+)GUG-nLuc-3XFLAG-CL1/PEST (Addgene plasmid #127317; a gift from Jeremy Wilusz^18^). The forward hPEST primer contained a 20 nucleotide tail complementary to the reverse primer used to amplify Renilla luciferase. The two PCR products, with a 20 nucleotide sequence overlap, were fused by PCR using the Renilla forward primer and the hPEST reverse primer, resulting in a Renilla-hPEST PCR product. The forward Renilla primer contained a tail with the XhoI restriction enzyme sequence and the hPEST reverse primer contained the NotI restriction enzyme sequence. This resulted in a final Renilla-hPEST PCR product flanked by XhoI and NotI restriction sequences that were used to replace eGFP in the original vector by standard cloning practices.

### Lentivirus production, transduction of A549 cells, and selection

293T cells were seeded in 6-well plates and transfected with the prISG15-Renilla-hPEST-IRES-Puro construct together with psPAX2 and pMD2.G (Addgene plasmids #12259 and #12260; gifts from Didier Trono) at a ratio of 2:1:1. FuGene HD (Promega; E2311) was used as the transfection reagent according to the manufacturer’s instructions. At 48h post-transfection, supernatants were harvested and filtered through a 0.45 µm syringe filter. Sub-confluent A549 cells were subsequently transduced with 2 mL of the filtered lentivirus-containing supernatant supplemented with 8 µg/mL polybrene (Sigma-Aldrich; 107689) for 24h. Transduced cells were then stimulated with 1000 IU/mL IFNα2 (Novusbio; NPB2-34971) for 4h to activate the *ISG15* promoter and thereby induce transcription of the monocistronic Renilla-hPEST-IRES-Puro transcript, which was followed by selection with puromycin (1 μg/mL, Thermo Fisher Scientific; A1113803) for 2 days. Cells were then subjected to limiting dilution in order to generate clonal cell populations.

### Luciferase assays

The 293T cell-based luciferase assay for detection of IFN-Is has been described previously^6,10^. For detection of IFN activity by A549-IFN-reporter (AIR) cells, 30,000 cells per well were seeded in a 96-well white-bottomed tissue culture plate and stimulated 24h later with IFNα2 (Novusbio; NPB2-34971), IFNβ (pbl assay science; 11420-1), IFNω (Novusbio; NBP2-35893), IFNγ (Novusbio; NBP2-34992), IFNλ1 (Novusbio; NBP2-34996), IFNλ2 (R&D systems; 8417-IL), or IFNλ3 (R&D systems; 5259-IL) at the indicated concentrations. Additionally, live-cell Renilla luciferase substrate (EnduRen, Promega; E6481) was added at a 1:10,000 dilution. At the indicated timepoints, luciferase signal was measured using a PerkinElmer EnVision plate reader (EV2104). Data were normalized to non-stimulated control cells, and expressed as relative Renilla activity in arbitrary units.

### Interferon neutralization assays

For detection of neutralizing anti-IFN antibodies, 1:100 diluted human plasmas and a dilution series of the indicated neutralizing anti-IFN antibodies (IFNβ, PBL assay science, 31410-1; IFNλ1: R&D systems, MAB15981; IFNλ2/3: R&D systems, MAB1587), or mock, were incubated with the appropriate IFN at the indicated concentration for 1h at room temperature with constant shaking at 600 rpm prior to addition to the appropriate reporter cell system (293T or AIR). For AIR cells, the medium was supplemented with EnduRen at a final dilution of 1:10,000. Cells were incubated for a further 24h at 37°C with 5% CO_2_ prior to determination of luciferase activities as described above.

## Abbreviations

AIR: A549 Interferon-Reporter
AutoAbs: autoantibodies
IFN: interferon
ISG: interferon-stimulated gene
IU: international units
prISG15: promoter of interferon-stimulated gene 15

## Data availability statement

The data that support the findings of this study are available from the corresponding author upon reasonable request.

## Acknowledgements

The authors thank Didier Trono and Jeremy Wilusz for gifting plasmids via Addgene, and the Swiss HIV Cohort Study (SHCS) for the use of anonymized human plasma samples. Financial support for this study was provided by the Academy of Medical Sciences (grant SBF003/1028 to DJH), the Wellcome Trust Institutional Strategic Support Fund (to DJH), and the Novartis Foundation for Medical-Biological Research (grant 23A069 to BGH).

